# Engineered placental organoids reveal molecular mechanisms underlying preeclampsia

**DOI:** 10.64898/2026.07.22.739976

**Authors:** Anya L. Arthurs, Caleb Lushington, Dulce L. Medina Garcia, Allanah Merriman, German A. Mora-Roldan, Laura Parry, Aafreen Boparai, Jose M. Polo, Fatwa Adikusuma, Paul Q. Thomas, Claire T. Roberts

## Abstract

Preeclampsia is a major pregnancy complication driven by placental dysfunction, yet research is limited by reliance on late stage patient-derived tissues and models that do not fully capture human disease. Here, we develop a genetically engineered human trophoblast organoid model of preeclampsia that can be generated without access to placental tissue. Using a CRISPR-based Prime Integrase strategy, we engineered induced trophoblast stem cells to constitutively express the preeclampsia-associated soluble fms-like tyrosine kinase-1 (sFlt-1) exon 15a isoform. Engineered organoids showed a transcriptional shift towards primary preeclamptic placentae and developed several features of disease. These included reduced PlGF, increased IL-6 and soluble endoglin, oxidative stress, impaired growth and an elevated sFlt-1/PlGF ratio comparable to primary preeclamptic trophoblast organoids. Importantly, these broader changes were not reproduced by adding recombinant human sFlt-1 to control organoids. Conditioned media from engineered organoids impaired endothelial network formation in both human umbilical vein and coronary artery endothelial cells, comparable to conditioned media from primary preeclamptic organoids. Treatment with either sulfasalazine or metformin normalised the sFlt-1/PlGF ratio and rescued impaired organoid growth. Together, these findings establish a scalable human model that links sustained expression of a disease-associated sFlt-1 isoform to trophoblast dysfunction and downstream endothelial dysfunction, providing a platform for mechanistic studies and drug discovery in preeclampsia.

## Introduction

Preeclampsia is a common and potentially life-threatening pregnancy complication characterised by maternal hypertension and multi-organ dysfunction [2], driven by placental disruption of angiogenic signalling. Central to its pathophysiology is excess production of soluble fms-like tyrosine kinase-1 (sFlt-1), an anti-angiogenic factor that antagonises vascular endothelial growth factor (VEGF) and placental growth factor (PlGF), leading to systemic endothelial dysfunction [3]. Elevated circulating sFlt-1, particularly an increased sFlt-1/PlGF ratio, is both a defining biomarker and a causal mediator of disease [4, 5].

Alternative splicing of *FLT1* generates multiple sFlt-1 isoforms, including variants incorporating exon 15a that are consistently elevated in preeclampsia and associated with disease severity [6]. Despite this, the functional consequences of exon 15a-containing sFlt-1 within human trophoblasts remain poorly defined. This is due in part to limitations of existing experimental models [7, 8]: patient tissues are restricted to late-stage disease and rely on access to primary tissue; animal models incompletely capture human placental biology; and *in vitro* trophoblast systems lack approaches to selectively manipulate endogenous transcript isoforms. Current genetic strategies in trophoblast models rely largely on transient overexpression or gene disruption, which do not recapitulate native transcript architecture or regulatory context. As a result, it has not been possible to determine whether expression of the exon 15a-containing isoform is sufficient to drive key features of trophoblast dysfunction.

Here, we address this gap by engineering selective expression of the exon 15a-containing sFlt-1 isoform in induced trophoblast stem cells (iTSCs), establishing a tractable human model of preeclampsia that does not rely on access to diseased placental tissue. Using a CRISPR-based Prime Integrase system, we integrated a constitutively expressed sFlt-1 exon 15a cassette at the AAVS1 safe-harbour locus, enabling selective and sustained expression of the disease-associated isoform and direct integration of its effects on trophoblast biology. We show that expression of sFlt-1 exon 15a is sufficient to alter trophoblast secretory profiles and activate cellular stress pathways, linking a clinically observed splicing event to functional disruption in human placental cells. These findings confirm the causal mechanistic role for exon 15a-containing sFlt-1 in trophoblast dysfunction and provide an experimental tool to refine understanding and accelerate drug discovery research in the preeclampsia field.

## Methods

### Cell Culture

Induced trophoblast stem cells [9] were kindly gifted by Professor Jose Polo. These cells, reprogrammed from adult human fibroblasts into trophoblast stem cells, are genetically very similar to early gestation primary trophoblast stem cells (also verified by our own analyses; see **Fig. 2a**). Cryopreserved cells were thawed at 37°C prior to resuspension in specialised trophoblast media as per Okae, *et al*. [10] and centrifugation (400 xg, 3 mins). Supernatant was removed and cells were resuspended prior to being seeded onto collagen IV-coated culture plates (5μg/mL; Sigma-Aldrich #C5533). Cells were cultured in a 37°C, 1% O_2_ and 5% CO_2_ incubator for 2 passages prior to transfection.

### Generation of plasmids

To generate the modified *sFlt-1* e15a overexpression organoid models, a prime editing-assisted site-specific integrase (eePASSIGE) strategy was employed, beginning with paired pegRNA-mediated prime editing (PE) integration of AttB at the *AAVS1* safe harbour locus. Briefly, a PEA1-Puro plasmid (Addgene #171991) was modified to replace the sgRNA domain with a second pegRNA cassette to permit dual-pegRNA activity from the single plasmid [11, 12]. Targeted construct assembly was achieved through *Bbs*I-mediated golden gate assembly of phosphorylated and annealed oligonucleotide sequences (Thermo Fisher), following optimised protocols for PEA1 assembly [12, 13]. This included *AAVS1* targeting guides and complementary AttB insertion templates with tevopreQ1 epegRNA motifs to direct donor integration. Donor plasmids were designed with an AttP recognition site for Bxb1 recombination with the *AAVS1* AttB locus, facilitated by pCMV-Bxb1 (eePASSIGE variant; Addgene #182142) [14], and an *sFLT-1* e15a splice variant cassette under the control of a CMV promoter. To ensure exclusive expression of the e15a variant, and not other splice variants, the expression cassette was designed around the 2kb *sFLT-1* e15a cDNA. This transcript underwent codon optimisation to simplify the synthesis of highly repetitive regions, and was followed by a bGH polyadenylation signal to ensure transcriptional termination. These sequences were synthesised as gBlocks (Twist Bioscience) and cloned into a pOG44 backbone via Gibson assembly. Subsequent plasmid purification was performed using the QIAprep Spin Miniprep Kit (Qiagen), and validation of oligonucleotide and gBlock insertion was carried out through Sanger sequencing (Australian Genome Research Facility [AGRF]). Oligonucleotide sequences can be found in **Table 1**.

**Table 1.**
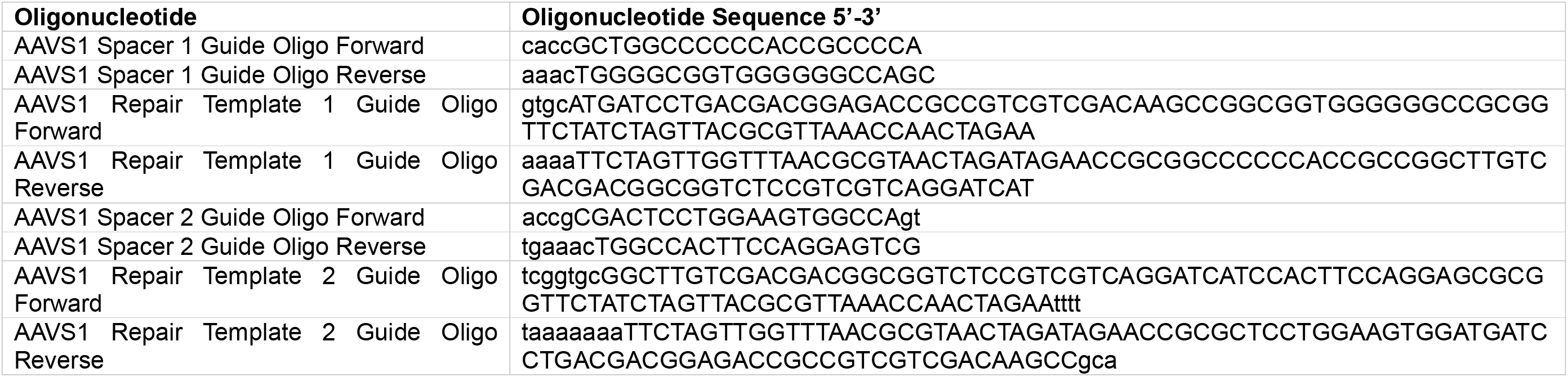
Oligonucleotide sequences for plasmid constructs.

### Induced trophoblast stem cell genetic modification

Induced trophoblast stem cells were transfected using the Neon™ Transfection System (Thermo Fisher Scientific), as indicated (*Neon Transfection User Guide, PubMAN0001557*). Prior to harvest for transfection, induced trophoblast stem cells were grown in antibiotic-free media for 1 passage. When cells reached ∼80% confluence, they were washed with PBS without calcium or magnesium (Gibco #14190144) and trypsinised (TryPLE Express; Gibco #12604013) then resuspended at a concentration of 1x10^7^ cells/mL. Cells were then washed with PBS without calcium or magnesium, prior to centrifugation (400 xg for 5 mins, room temperature). PBS was aspirated and the cell pellet was resuspended in the Neon™ Resuspension Buffer R. 100 μL of the resultant cell suspension was mixed with 10 μg total plasmid DNA (where Donor:PE:BXBl plasmids were transfected at a 1:1:1 ratio) before being loaded into the Neon™ Pipette. Transfection parameters were 1600 V, 20 ms, 2 pulses. After electroporation, cells were immediately seeded into collagen IV-coated culture plates containing prewarmed trophoblast media without antibiotics.

After 24 h, puromycin (0.2μg/mL) was added to the culture media to begin antibiotic selection of successfully transfected cells (as previous, [15]). Puromycin selection continued for 14 days of culture (until cells reached confluence) prior to junction PCR and Sanger sequencing to confirm plasmid integration.

### Tissue collection

For primary organoids: First-trimester human placentae (6–7 weeks’ gestation) were collected with informed consent from women undergoing elective termination of pregnancy at the Pregnancy Advisory Centre, Queen Elizabeth Hospital, Woodville, South Australia. Ethical approval was granted by the Central Adelaide Local Health Network Human Research Ethics Committee (HREC/16/TQEH/33; Q20160305). Placentae were collected within minutes of the procedure, after which villous tissue was washed in Hanks’ Balanced Salt Solution (HBSS; Gibco, Sigma-Aldrich) and transported to the laboratory on ice.

Term placentae for sequencing: As previously described[16], term placentae were collected at the Lyell McEwin Hospital, Elizabeth, South Australia, from women enrolled in the Screening Tests to predict poor Outcomes of Pregnancy (STOP; 2015–2018) cohort study[17]. The study was registered with the Australian and New Zealand Clinical Trials Registry (STOP: ACTRN 12614000985684).

Placental tissue biopsies were washed in phosphate-buffered saline (PBS), immersed in RNAlater Stabilisation Solution (Thermo Fisher Scientific, Massachusetts, USA), and stored at −80°C. Ethical approval was granted by the Women’s and Children’s Health Network Human Research Ethics Committee (HREC/14/WCHN/90). Written informed consent was obtained from all participants.

For term preeclamptic organoids: Term preeclamptic placentae were obtained with informed consent after delivery from Flinders Medical Centre, in Bedford Park, South Australia. Ethics were obtained from the Southern Adelaide Local Health Network Human Research Ethics Committee (2024/HRE00218).

Late onset preeclampsia was defined as the onset of preeclampsia after 34 weeks’ gestation. Preeclampsia was defined as (peripheral) hypertension [systolic BP (SBP) ≥ 140 mmHg or diastolic BP (DBP) ≥ 90 mmHg] after 20 weeks of gestation in previously normotensive women, with proteinuria (24 h urinary protein ≥ 300 mg or spot urine protein: creatinine ratio ≥ 30 mg/mmol creatinine or urine dipstick protein ≥ 2+) or any multi-organ complication of preeclampsia [17]. The estimated date of delivery was calculated from a certain last menstrual period (LMP) date and was only adjusted if either (1) a scan performed at < 16 weeks of gestation found a difference of ≥7 days between the scan gestation and that calculated by the LMP or (2) on 20-week scan a difference of ≥ 10 days was found between the scan gestation and that calculated from the LMP. If the LMP date was uncertain, then scan dates were used to calculate the estimated date of delivery. The clinical characteristics of all the participants are summarized in **Supplementary Table 1**. No significant differences were observed between term healthy and preeclampsia groups (n = 3 healthy, n = 6 preeclampsia). A subset of preeclampsia (n = 3) was sequenced while all preeclampsia tissues were used for organoid generation.

### Isolation of TSCs

Primary organoids were formed from early gestation tissue as per Arthurs *et al*.[15]. Briefly, TSCs were isolated from first-trimester placentae using the method described by Dietrich *et al*. [18]. Villous tissue was washed in HBSS (4 °C), and individual villous structures were manually dissected for subsequent processing. Villi suspended in HBSS were centrifuged at 1000 rpm for 1 min, followed by three sequential enzymatic digestion steps and density gradient centrifugation according to Haider *et al*. [19]. Cells located at the interface between the 35% and 50% Percoll layers were carefully collected, thoroughly washed in HBSS, and seeded onto fibronectin-coated culture plates in stemness-promoting medium. TSC medium consisted of DMEM/F12 (Gibco) supplemented with 1× B-27 (Gibco), 1× Insulin-Transferrin-Selenium-Ethanolamine (ITS-X; Gibco), 1 µM A83-01 (Tocris), 50 ng/mL recombinant human epidermal growth factor (rhEGF; R&D Systems), 2 µM CHIR99021 (Tocris), and 5 µM Y27632 (Santa Cruz).

Trophoblast organoids were generated from full term (40 weeks’ gestation) preeclamptic placentae as previously described [20]. Briefly, placental villi were dissected and washed in HAMS/F12 (#11765054, Gibco) prior to dissociation using a 0.2% Trypsin / 0.02% EDTA solution. After filtration, undigested tissue was further processed using collagenase V. Pooled isolated cells were centrifuged (400 xg, 5 min) and washed (Advanced DMEM/F12 medium, #12634010, Gibco).

### Organoid culture

Organoids created from both iTSCs and primary TSCs were created following the same protocol. Instead of embedding in a synthetic extracellular matrix, trophoblast stem cells were seeded at 1×10^5^ cells per well with 3 mL total organoid media (per Yang, et al. with suggested media additions[21]) in a Corning® Costar® Ultra-Low Attachment Multiple Well Plate (#CLS3471, Merck) and placed on an orbital shaker (85 rpm, Thermo Scientific™, Cat. 88881102, orbit 1.9 cm) in an incubator (37 °C, 5% O_2_, 5% CO_2_).

The Incucyte® SX5 Live-Cell Analysis System (Sartorius) system was used to assess total organoid area, using the Organoid Software program (Sartorius) in-built settings for analysis.

### Treatment compounds

To assess the effect of recombinant human sFlt-1 addition to culture media, control organoids were supplemented with Recombinant Human Soluble Fms-Like Tyrosine Kinase-1 (Creative Diagnostics, Düsseldorf, Germany; #DAG-WT128) at a final concentration of 1800 pg/mL in culture medium. Recombinant sFlt-1 was replenished with each media change for the duration of culture.

To assess drug effectiveness, organoids were treated with one of the following compounds for a duration of 7 days:

1. Sulfasalazine (Sigma, #S0883) at a final concentration of 200 µM in culture medium. Dosage adapted from Brownfoot, *et al*.[22]

2. Metformin (Sigma, #317240) at a final concentration of 1 mM in culture medium. Dosage adapted from Brownfoot, *et al*.[23]

### Conditioned media harvesting and processing

Following establishment, culture medium was removed from control, sFlt-1 OEx and primary preeclamptic trophoblast organoids and the organoids were gently washed with PBS. Fresh organoid medium, as described above but with FBS replaced with Knockout™ Serum Replacement (Gibco #10828028), was added and organoids were cultured for 24 h. Conditioned media was then collected and centrifuged (400 ×g, 5 mins) to remove detached cells and cellular debris. The resulting supernatant was passed through a 0.45 μm filter before use in downstream culture assays.

### Culture Assays

Primary Human Umbilical Vein Endothelial Cells (HUVEC): cells were obtained from ATCC and grown using the Endothelial Cell Growth Kit-VEGF (ATCC, #PCS-100-041) according to manufacturer’s instructions. Once cells reached 80% confluence in a 25 cm^2^ flask, they were moved to a 75 cm^2^ flask. At 60% confluence, the culture media was changed to incorporate 0.2% FBS instead of the recommended 2% FBS, to serum starve cells for 24 h prior to assay.

After 24 h of serum starvation, a 96-well plate was pre-coated with a synthetic hydrogel matrix (1:1 Vitrogel® Hydrogel Matrix to culture medium; 50 μL total). HUVEC cells were seeded at 1×10^4^ cells/well in 100 μL of culture medium containing either:

1. Normal cell culture media supplemented with 50 ng/mL VEGF.
2. Normal cell culture media (Endothelial Cell Growth Kit-VEGF; ATCC, #PCS-100-041);
3. 1:50 ratio of primary preeclamptic organoid conditioned media to normal cell culture media;
4. 1:50 ratio of sFlt-1 OEx organoid conditioned media to normal cell culture media;
5. 1:100 ratio of sFlt-1 OEx organoid conditioned media to normal cell culture media;
6. 1:500 ratio of sFlt-1 OEx organoid conditioned media to normal cell culture media; or
7. 1:1000 ratio of sFlt-1 OEx organoid conditioned media to normal cell culture media.

The Incucyte® SX5 Live-Cell Analysis System (Sartorius) system was then used to monitor tube formation, using the Angiogenesis Analysis Software program (Sartorius).

Primary Human Coronary Artery Endothelial Cells (HCAEC): cells were obtained from ATCC and grown using the Endothelial Cell Growth Kit-VEGF (ATCC, #PCS-100-041) according to manufacturer’s instructions. Once cells reached 80% confluence in a 25 cm^2^ flask, they were moved to a 75 cm^2^ flask. At 60% confluence, the culture media was changed to incorporate 0.2% FBS instead of the recommended 2% FBS, to serum starve cells for 24 h prior to assay.

After 24 h of serum starvation, a 96-well plate was pre-coated with a synthetic hydrogel matrix (1:1 Vitrogel® Hydrogel Matrix to culture medium; 50 μL total). HCAEC cells were seeded at 1×10^4^ cells/well in 100 μL of culture medium containing either:

1. Normal cell culture media supplemented with 50 ng/mL VEGF.
2. Normal cell culture media (Endothelial Cell Growth Kit-VEGF; ATCC, #PCS-100-041);
3. 1:50 ratio of primary preeclamptic organoid conditioned media to normal cell culture media;
4. 1:50 ratio of sFlt-1 OEx organoid conditioned media to normal cell culture media;
5. 1:100 ratio of sFlt-1 OEx organoid conditioned media to normal cell culture media;
6. 1:500 ratio of sFlt-1 OEx organoid conditioned media to normal cell culture media; or
7. 1:1000 ratio of sFlt-1 OEx organoid conditioned media to normal cell culture media.

Endothelial network formation was imaged for 6 h using the Incucyte® SX5 Live-Cell Analysis System. Network formation was quantified at 6 h using Incucyte Angiogenesis Analysis Software, with number of branching points and mean mesh diameter used as outcome measures. Five images per well were analysed, with all conditions performed in biological triplicate as well as technical triplicate (final n=9).

### RNA Extraction

Culture media: Culture medium was removed from organoid cultures. Samples were mixed with 600 µL Buffer RLT Plus (RNeasy Plus Mini Kit; QIAGEN, Victoria, Australia). Total RNA was subsequently isolated using the RNeasy Plus Mini Kit (QIAGEN) in accordance with the manufacturer’s instructions. RNA quantity and quality were assessed using a NanoDrop spectrophotometer (Thermo Fisher Scientific), and only samples with A260/A280 and A260/A230 ratios >1.9 were included for downstream analyses.

Tissue: Term placental tissue (0.025 g) was weighed and washed in phosphate-buffered saline (PBS). Samples were homogenised in 600 μL Buffer RLT Plus (RNeasy Plus Mini Kit; QIAGEN, Victoria, Australia) for 3.5 min at 30 Hz using a TissueLyser (QIAGEN). Total RNA was subsequently isolated from the resulting supernatant using the RNeasy Plus Mini Kit (QIAGEN) in accordance with the manufacturer’s instructions. RNA purity and integrity were assessed using the Experion™ system (Bio-Rad, New South Wales, Australia), and only samples with an RNA integrity number (RIN) ≥8 were used.

Organoids: Organoids were collected and washed in PBS before the addition of 600 μL Buffer RLT Plus (RNeasy Plus Mini Kit; QIAGEN, Victoria, Australia). Samples were incubated for 20 min, after which total RNA was extracted using the RNeasy Plus Mini Kit (QIAGEN) according to the manufacturer’s instructions. RNA purity and integrity were assessed using the Experion™ system (Bio-Rad, New South Wales, Australia), and only samples with an RNA integrity number (RIN) ≥8 were used.

### Junction PCR

Puromycin-selected transfected cells were checked for correct plasmid integration by PCR, using primers which overlapped junctions 1 and 2 (as indicated in **Fig. 1d**; primer sequences included in **Table 2**). 100 ng of gDNA was used for PCR, using the Q5® High-Fidelity PCR Kit (New England Biolabs, #E0555L) according to the manufacturer’s instructions. After initial denaturation at 98 °C for 30 s, 35 cycles of denaturation (98 °C, 10 s), annealing (65 °C, 20 s) and extension (72 °C, 20 s) were completed. The resultant PCR products were then electrophoresed through a 1.8% agarose gel for 1 h at 100V and bands were compared to a standardised ladder (NEB 50bp ladder, #N3236S). PCR products spanning junctions 1 and 2 were purified using the Monarch® Spin PCR & DNA Cleanup Kit (NEB #T1130S) and subjected to Sanger sequencing by the Australian Genome Research Facility (AGRF). Resulting sequences were aligned against the predicted AAVS1-donor junction sequences using SnapGene software.

**Figure 1.**
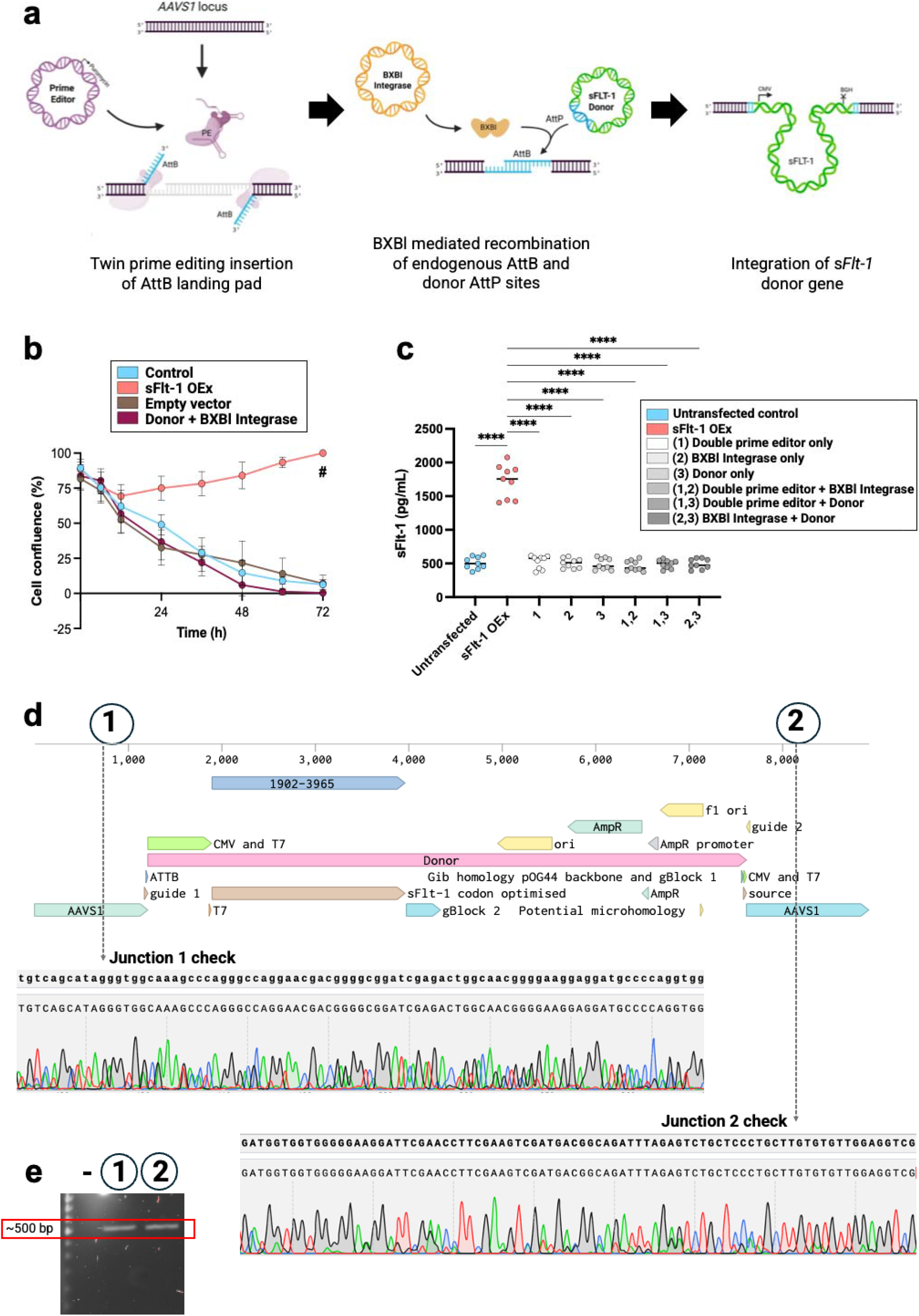
Transfection strategy and checks. **a,** schematic of the CRISPR Prime Integrase approach used for targeted integration of the sFLT-1 gDNA with constitutive overexpression at a safe harbour. This approach involved use of a double prime editor, along with BXBI integrase and donor plasmid. sFlt-1 exon 15a isoform was confirmed for overexpression in induced trophoblast stem cells (iTSCs). **b,** Confluence (%) of cells grown for 72 hours in media containing 0.2μg/mL puromycin to select for successful transfection. Control (blue) cells were untransfected; sFlt-1 OEx (coral) cells were transfected with all three plasmids (double prime editor, BXBl integrase and donor plasmid); empty plasmid (brown) cells were transfected with a commercially available empty plasmid backbone; donor + BXBl integrase (maroon) cells were transfected with only two of the three plasmids (BXBl integrase and donor plasmid only). **c,** Concentration of sFlt-1 (pg/mL) in culture media of cells transfected with different combinations of the three plasmids, where only “sFlt-1 OEx” (coral) integrated all three plasmids successfully. **d,** Genomic map of AAVS1 after donor plasmid integration. Numbers 1 and 2 indicate junctions between the sequences which checked in transfected cells using Sanger sequencing. Sanger sequencing confirmed perfect sequence complementary to **(1)** Junction 1 and **(2)** Junction 2 between the expected sequence (top) and observed in sFlt-1 OEx cells (bottom). **e,** PCR product size for junction 1 and junction 2 checks. Negative control indicated with **–** sign. # and **** indicate statistical significance (p < 0.0001), as determined by one-way ANOVA with multiple comparisons.

**Figure 2.**
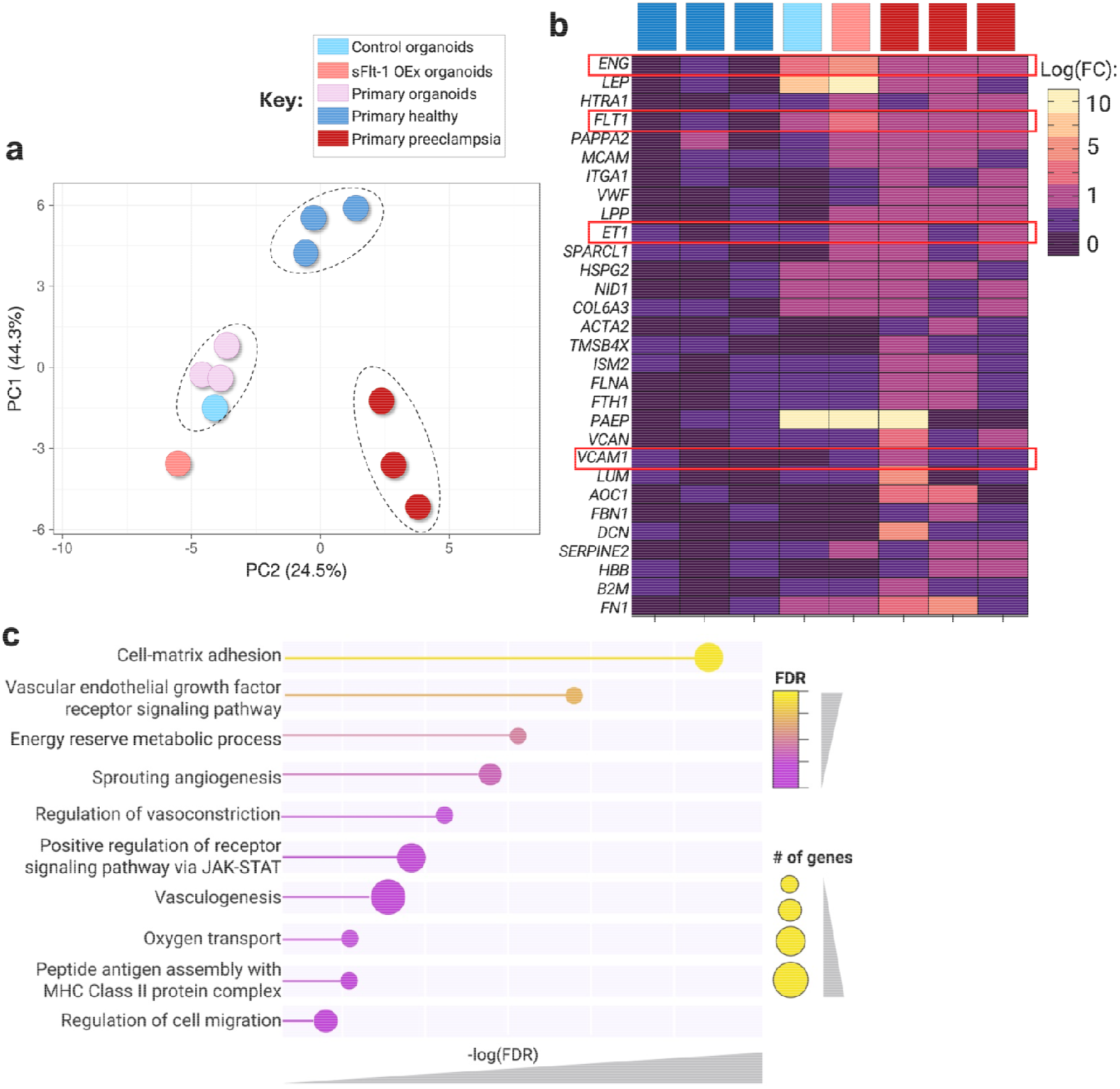
sFlt-1 overexpression trophoblast organoids recapitulate transcriptional features of preeclampsia. **a,** Principal component analysis of global transcriptomic profiles from control organoids (blue), sFlt-1 exon 15a overexpression (sFlt-1 OEx) organoids (coral), primary early gestation organoids (pink), and primary placenta samples (healthy [red] and preeclampsia [navy]). **b,** Heatmap of top differentially expressed genes across experimental groups, showing log fold change (logFC) in expression compared with healthy controls. Key genes associated with preeclampsia pathology are indicated by red boxes. **c,** Gene ontology enrichment analysis of differentially expressed genes in sFlt-1 OEx organoids relative to control organoids. Colour of the bubble indicates false discovery rate (FDR) value. Size of the bubble indicates the number of genes out of the top differentially expressed genes pertaining to that gene ontology term. -logFDR value is indicated by the length of the line across the x-axis. Sequencing experiments included n=3 patient samples per primary tissue group. Source data file is provided.

**Table 2.**
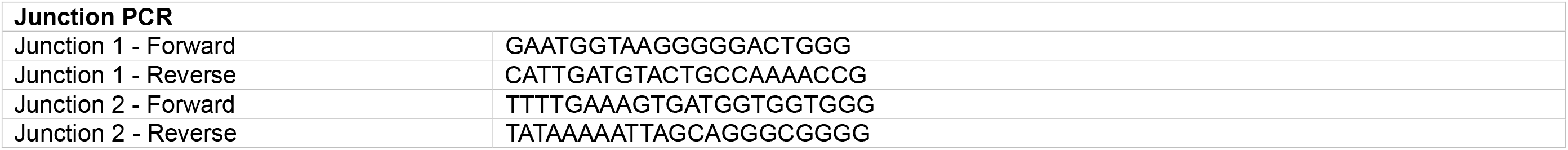
Sequences of primers used.

### Sequencing and Bioinformatic Analyses

Library preparation and High Throughput Sequencing was performed by the South Australian Genomics Facility (SAGC) using a NovaSeq X Illumina sequencer. All bioinformatic support, including alignment and quality control metrics, were performed by SAGC as part of service. Briefly, differential gene expression analysis was performed in R (v4.0.5) using the *edgeR* (v3.16.5; [24]) and *limma* (v3.30.11; [25]) packages. Initially, lowly expressed mRNAs were removed and normalisation for library composition bias was undertaken using *edgeR*. Expression data were subsequently normalised using the Trimmed Mean of M-values (TMM) method. The *limma* package was then applied to generate sample weights and perform log transformation, with the voom function used to model the mean–variance relationship across observations and incorporate these estimates into the normalised log-count data. Differential expression testing, including calculation of moderated F-statistics, adjusted *P*-values, and log fold changes, was carried out using a moderated *t*-test [26]. Multiple testing correction was performed using the Benjamini–Hochberg method [27], and genes with a false discovery rate (FDR) ≤ 0.05 were considered significantly differentially expressed. The gene ontology analysis was generated through the use of QIAGEN IPA (QIAGEN Inc., https://digitalinsights.qiagen.com/IPA [28]).

### Enzyme-linked ImmunoSorbent Assay (ELISA)

As previous [20], angiogenic factors sFlt-1 and PlGF were quantified in trophoblast organoid culture media. Measurements were performed according to the manufacturers’ protocols using the PikoKine® sFlt-1 ELISA kit (#MBS175839) and the Quantikine Human PlGF ELISA kit (#DPG00), respectively. An sFlt-1 exon 15a-specific ELISA was established as previously by Palmer, *et al*. [6, 29]. IL-6, TNF-α and soluble endoglin expression in trophoblast organoid culture media were measured using the Sigma-Aldrich Human IL-6 ELISA kit (#EZIL6), Millipore Human TNF-alpha ELISA kit (#EZHTNFA-150K) and Invitrogen Human Endoglin (CD105) ELISA kit (#EHENG), respectively.

### Immunofluorescence

Immunofluorescence staining was performed using a previously described protocol [30], with minor modifications. Organoids were washed in phosphate-buffered saline (PBS) and fixed in 4% paraformaldehyde for 20 min at room temperature. Samples were permeabilised for 15 min in PBS containing 0.3% Triton X-100 and 3% bovine serum albumin (BSA), followed by blocking in 3% BSA in PBS for 30 min. Samples were then incubated overnight at 4 °C with primary antibodies against PSG1 (Anti-PSG1 Rabbit Polyclonal Primary Antibody, Abcam #ab233130, 1:200) and 8-OHdG (Anti-8-OHdG (E-8) Mouse Monoclonal Antibody, Santa Cruz #sc-393871, 1:100) diluted in PBS containing 0.1% BSA. Following three washes in PBS containing 0.1% Tween-20, samples were incubated with Alexa Fluor-conjugated secondary antibodies against rabbit IgG (Alexa Fluor 488 Goat Anti-Rabbit IgG H&L, Abcam #ab150077, 1:500) and mouse IgG (Alexa Fluor 647 Goat Anti-Mouse IgG H&L, Thermo Fisher #A32728, 1:500) for 30 min at room temperature. Samples were washed a further three times before nuclei were counterstained with NucBlue™ Fixed Cell ReadyProbes™ Reagent (DAPI) for 10 min. Samples were subsequently mounted in 15 µL Fluoromount-G mounting medium (Invitrogen, #00-4958-02). Imaging was performed using an Olympus FV3000 confocal microscope, and images were analysed using FV31S-SW FLUOVIEW software.

### Statistics and Reproducibility

All statistical analyses were performed using SPSS software and graphs were generated using GraphPad Prism. Data are presented as mean ± standard deviation unless otherwise stated. For all experiments, *p* < 0.05 was considered significant.

All ELISAs were completed on n = 3 biological replicates per group, in technical triplicate (total n=9).

Comparisons between two groups (e.g., control vs sFlt-1 OEx) were analysed by two-tailed Mann–Whitney tests. For multiple-group comparisons, Welch’s ANOVA with post hoc multiple-comparison correction was applied. For time course comparisons, a repeated-measures two-way ANOVA was used. Exact statistical tests and sample sizes are indicated in figure legends. No statistical method was used to predetermine sample size. All experiments were independently reproduced at least three times with consistent results.

## Results

### Establishment of an in vitro preeclamptic trophoblast organoid model

We first sought to create an *in vitro* trophoblast organoid model of preeclampsia that does not rely on access to human primary tissue but instead recapitulates key molecular features of the condition. To achieve this, we focused on overexpression of the exon 15a-containing variant of *FLT1*, which is consistently elevated in the preeclamptic placenta and contributes to excess production of soluble fms-like tyrosine kinase-1 (sFlt-1).

To enable selective expression of this isoform, we developed a CRISPR-based Prime Integrase strategy to introduce a constitutively expressed sFlt1 exon 15a cassette into the AAVS1 safe-harbour locus of induced trophoblast stem cells (iTSCs) (**Fig. 1a**). This approach combines twin prime editing to install a recombinase specific genomic landing pad at the AAVS1 locus, followed by site-specific recombination to integrate an exon 15a-containing sFlt-1 donor with CMV promoter construct [31]. The donor construct contained the sFlt-1 exon 15a coding sequence, enabling selective and sustained expression of the disease-associated isoform.

To generate the sFlt-1 exon 15a overexpression line, transfected cells were first subjected to puromycin selection, resulting in selective survival of cells receiving the prime editing system (**Fig. 1b**). To confirm that sFlt-1 overexpression required the complete editing system, cells were transfected with different combinations of the double prime editor, BXBl integrase and donor plasmids. Increased sFlt-1 concentrations in culture media were observed only when all three plasmids were delivered together, with no increase detected in the control of incomplete plasmid combinations (**Fig. 1c**). The AAVS1 integration site and predicted donor junctions used to validate successful sFlt1 e15a cDNA insertion are shown in **Fig. 1d**. Targeted integration of the donor construct at AAVS1 was confirmed by 500bp PCRs (**Fig. 1e**) over both the 5’ and 3’ integration boundaries (labelled **1** and **2**). Further validation of the correct donor integration was performed by Sanger sequencing at the donor junctions. Together, these analyses confirmed accurate targeted insertion of the constitutively expressed sFlt-1 exon 15a cassette at the AAVS1 locus.

Following validation, engineered iTSCs were differentiated into trophoblast organoids for downstream analyses.

### sFlt-1 Overexpression trophoblast organoids recapitulate gene expression signatures of preeclamptic placenta

To determine the transcriptional impact of exon 15a-containing sFlt-1 overexpression, we performed RNA sequencing on:

- Control organoids: generated from iTSCs
- sFlt-1 overexpression organoids: iTSCs underwent genetic editing prior to organoid formation
- Primary organoids: organoids generated from trophoblast stem cells isolated from 6 weeks’ gestation placentae (n=3)
- Primary healthy tissue: Term placenta tissue from patients with uncomplicated pregnancies (n=3)
- Primary preeclampsia tissue: Term placenta tissue from patients with late-onset preeclampsia (n=3)

To assess global transcriptional changes, we performed principal component analysis on RNA-sequencing data (**Fig. 2a**). Samples segregated along principal component 1 (PC1; 44.3% variance) and principal component 2 (PC2; 24.5% variance), with clear clustering by sample type. Control organoids and primary organoids (early gestation) formed a distinct cluster, indicating the biological recapitulation of the early gestation placental transcriptome by the control organoid (generated from iTSCs). sFlt-1 overexpression organoids separated from control and occupied an intermediate position between control organoid and primary preeclampsia samples. Notably, primary preeclamptic samples clustered separately from the primary healthy placenta samples, and sFlt-1 overexpression organoids shifted toward the transcriptional space defined by preeclamptic samples. These data indicate that induction of the exon 15a-containing sFlt-1 isoform is associated with broad transcriptional changes that partially recapitulate the molecular profile of the preeclamptic placenta.

To assess the extent to which sFlt-1 overexpression organoids reproduce disease-associated transcriptional signatures, we compared expression of top differentially expressed genes between control and sFlt-1 overexpression organoids, and included expression of these selected genes across primary placental samples. Hierarchical clustering demonstrated that sFlt-1 overexpression organoids more closely resembled primary preeclamptic samples than did the control organoids (**Fig. 2b**).

30 genes were differentially expressed in sFlt-1 overexpression organoids, compared to controls, as a consequence of sFlt-1 exon 15a overexpression. Notably, expression of key angiogenic regulators including *FLT1* and endoglin *(ENG)*, vasoconstrictor endothelin 1 (*ET1*) and inflammatory vascular cell adhesion molecule 1 (*VCAM1*) were spontaneously upregulated in sFlt-1 overexpression organoids.

To identify biological processes altered by sFlt-1 overexpression, we performed Gene Ontology Biological Process enrichment analysis on differentially expressed genes between control and sFlt-1 overexpression organoids (**Fig. 2c**). The top ten processes (as determined by -logFDR) are depicted. This revealed significant enrichment of processes related to cell matrix adhesion (*-logFDR*=3.68; 6 genes), vascular endothelial growth factor signalling (*-logFDR*=2.30; 1 gene), energy reserve metabolic processes (*-logFDR*=2.10; 2 genes), sprouting angiogenesis (*-logFDR*=1.92; 4 genes) and regulation of vasoconstriction (*-logFDR*=1.74; 3 genes). Additional enriched terms included positive regulation of receptor signalling pathway via JAK-STAT (*-logFDR*=1.70; 6 genes), vasculogenesis (*-logFDR*=1.66; 8 genes), oxygen transport (*-logFDR*=1.57; 2 genes), peptide antigen assembly with MHC Class II protein complex (*-logFDR*=1.56; 1 gene) and regulation of cell migration (*-logFDR*=1.40; 4 genes). Together, these data indicate coordinated changes in biological processes central to placental vascular function and cellular stress responses.

### sFlt-1 overexpression organoids independently recapitulate several pathological features of preeclampsia

Given that sFlt-1 overexpression was the primary goal of our genetic edits, we first sought to confirm this. Across 4 weeks of culture, sFlt-1 overexpression organoids secreted significantly elevated concentrations of sFlt-1 into culture media, compared with controls (mean 1748.75 pg/mL versus 515.98 pg/mL; **Fig. 3a**). To determine whether the effects observed in our engineered model could be reproduced by extracellular exposure to sFlt-1 alone, control organoids were also cultured in the presence of recombinant human sFlt-1 ((rh) sFlt-1). As expected, addition of (rh) sFlt-1 significantly increased the concentration of total sFlt-1 detected in culture media to levels comparable to those produced by our sFlt-1 overexpression organoids (mean 1785.99 pg/mL versus 1748.75 pg/mL).

**Figure 3.**
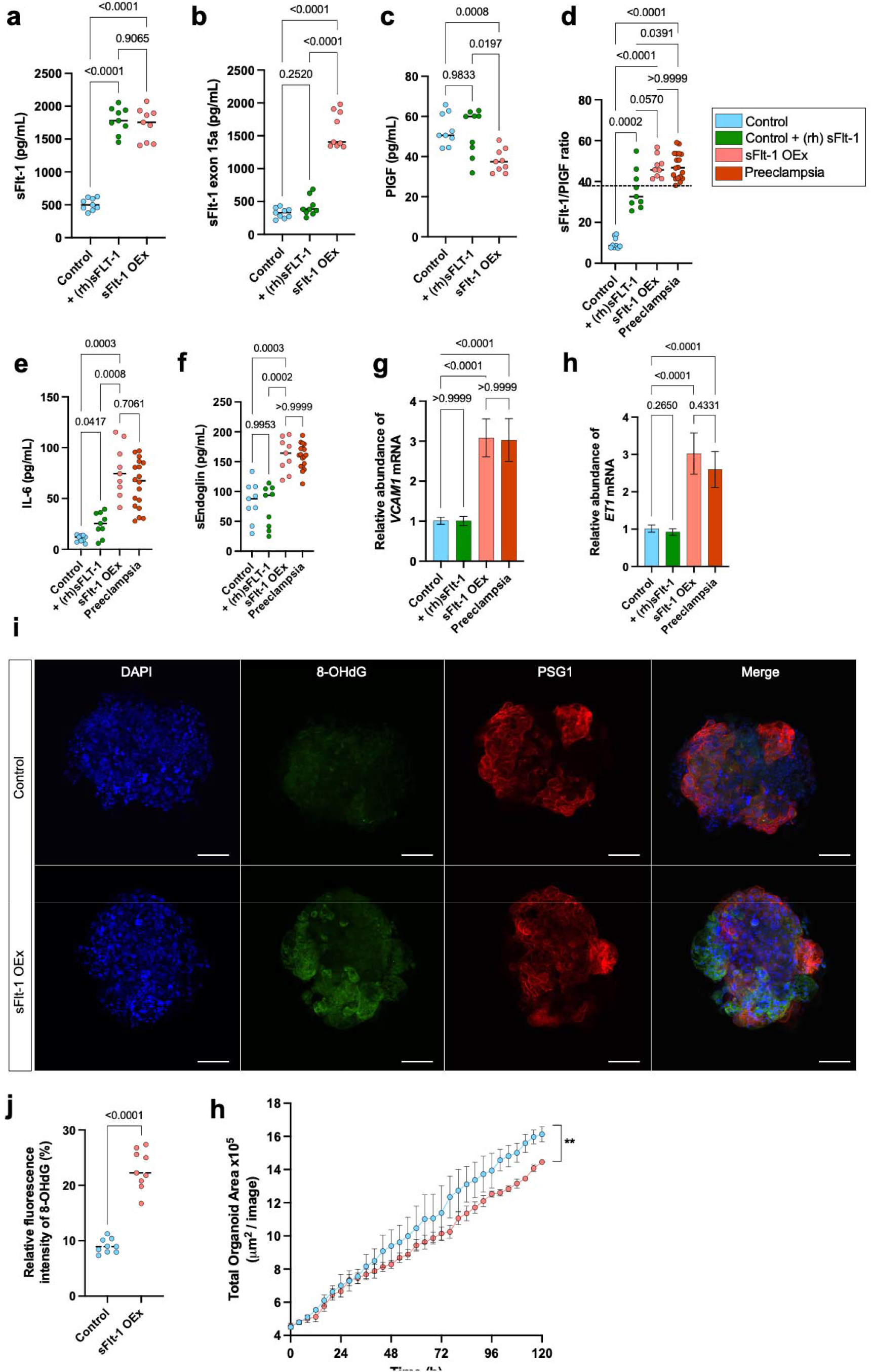
Molecular characterisation of sFlt-1 exon 15a overexpression trophoblast organoids. Concentrations of angiogenic factors in culture media of control (blue), control supplemented with (rh) sFlt-1 (green), sFlt-1 OEx (coral) and primary preeclamptic (red) trophoblast organoids, including **a,** sFlt-1 (pg/mL) **b,** sFlt-1 exon 15a (pg/mL), **c,** PlGF (pg/mL) **d,** sFlt-1/PlGF ratio. The dashed line indicates the clinical cutoff for preeclampsia diagnosis [1]. **e,** IL-6 (pg/mL) and **f,** sEndoglin (pg/mL) concentrations in organoid culture media. **g,** Relative abundance of VCAM1 mRNA and **h,** ET1 mRNA. **i,** Representative immunofluorescence images of sFlt-1 overexpression and control organoids stained for 8-OHdG (oxidative stress, green), PSG1 (syncytiotrophoblast, red) and DAPI (nuclei, blue). Scale bar, 100 μm. **j,** Quantification of 8-OHdG fluorescence intensity in sFlt-1 overexpression and control organoids. **k,** Organoid growth measured as total organoid area x10^5^ of control (blue) and sFlt-1 overexpression (coral) trophoblast organoids, using the Incucyte SX5 Live-Cell Imaging Platform with Incucyte Organoid Analysis Software. For **a-f**, data are presented as a scatter plot with median indicated with a horizontal line and assessed using Welch’s ANOVA with multiple comparisons. For **g-h,** data are presented as a histogram with mean ± standard deviation and assessed using Welch’s ANOVA with multiple comparisons. For **j,** data are presented as a scatter plot with median indicated with a horizontal line and assessed using a Mann-Whitney test. For **k**, data are presented as mean ± standard deviation and assessed using a two-way ANOVA. n=9 trophoblast organoid replicates per group for control, control + (rh) sFlt-1 and sFlt-1 OEx. n = 18 trophoblast organoid replicates per group for term preeclamptic organoids. *\*\*p < 0.01. Source data file is provided*.

We next sought to determine the proportion of sFlt-1 corresponding specifically to the exon 15a isoform. Consistent with the model design, concentrations of sFlt-1 exon 15a in culture media were significantly higher in sFlt-1 overexpression organoids compared with controls and controls + (rh) sFlt-1 (mean 1587.09 pg/mL versus 323.72 pg/mL and 421.38 pg/mL, respectively; **Fig. 3b**). Notably, sFlt-1 exon 15a concentrations closely mirrored total sFlt-1 levels, indicating that the majority of secreted sFlt-1 produced by the engineered organoids corresponded to the disease-associated exon 15a isoform.

Importantly, the engineered organoids spontaneously developed several additional pathological features of preeclampsia that were not fully recapitulated by direct addition of (rh) sFlt-1 to the culture media. PlGF, a pro-angiogenic molecule critical for feto-placental circulation and trophoblast growth [32], was significantly reduced in culture media of sFlt-1 overexpression organoids compared with both control and control + (rh) sFlt-1 organoids (mean 37.95 pg/mL versus 53.51 pg/mL and 52.01 pg/mL, respectively; **Fig. 3c**). This consequently altered the ratio of sFlt-1/PlGF as detected in culture media. The sFlt-1/PlGF ratio in maternal circulation is the most widely used clinical biomarker for preeclampsia, reflecting placental dysfunction [1]. A clinical cut-off of 38 or higher signifies elevated risk of preeclampsia (indicated in diagrams with a dotted line; **Fig. 3d**). The sFlt-1/PlGF ratio was significantly elevated in sFlt-1 overexpression organoids compared with controls (mean 47.15 versus 9.87). This ratio was also measured in the culture media of trophoblast organoids generated from primary term preeclamptic placentae [20], and was not significantly different from that observed in sFlt-1 overexpression organoids (mean 48.37 versus 47.15).

We next assessed secretion of key pro-inflammatory cytokines, namely interleukin 6 (IL-6) and tumour necrosis factor α (TNF-α), which are elevated in maternal circulation of preeclamptic pregnancies [33–35]. Whilst levels of TNF-α were below assay detection limit, IL-6 concentration in culture media was significantly elevated in sFlt-1 overexpression organoids compared with control and control + (rh) sFlt-1 organoids (mean 77.56 pg/mL versus 10.66 pg/mL and 24.60 pg/mL, respectively; **Fig. 3e**). Levels of IL-6 were not significantly different between sFlt-1 overexpression organoids and primary preeclamptic organoids (mean 77.56 pg/mL versus 64.12 pg/mL, respectively). We also measured secretion of soluble endoglin (sEndoglin) into culture media.

Like sFlt-1, soluble endoglin is an anti-angiogenic factor that is characteristically elevated in maternal circulation of preeclamptic pregnancies. Soluble endoglin concentrations were likewise elevated in the culture media from sFlt-1 overexpression organoids compared with controls (mean 160.26 pg/mL versus 72.81 pg/mL; **Fig. 3f**). Levels of soluble endoglin were not significantly different between sFlt-1 overexpression organoids and primary preeclamptic organoids (mean 160.26 pg/mL versus 159.10 pg/mL, respectively). Direct addition of (rh) sFlt-1 into culture media did not significantly alter soluble endoglin secretion (mean 74.68 pg/mL), further indicating that the broader anti-angiogenic phenotype of the engineered organoids cannot be reproduced by extracellular sFlt-1 exposure alone.

We also measured trophoblast organoid growth over time. Whilst term preeclampsia is not consistently associated with smaller placentae [36], low weight placentae are more commonly observed with preeclampsia [37]. Measurement of total organoid area over time demonstrated significantly impaired growth of sFlt-1 overexpression organoids compared with controls (**Fig. 3g**), indicating slower growth.

Finally, we examined oxidative stress in the trophoblast organoids. Placental oxidative stress is a well-characterised part of preeclampsia pathology [38]. Immunofluorescence revealed significantly increased 8-OHdG staining in sFlt-1 overexpression organoids compared with controls (relative fluorescence intensity mean 23.24% versus 8.93%; **Fig. 3h,i**). 8-OHdG staining was distributed throughout the organoids, indicating widespread oxidative stress, including in the syncytiotrophoblast (stained with PSG1).

### sFlt-1 overexpression trophoblast organoids respond to clinically used therapeutics for preeclampsia

To determine whether sFlt-1 overexpression trophoblast organoids recapitulate clinically relevant features of preeclampsia, we assessed their responsiveness to therapeutics reported to modulate angiogenic imbalance, including sulfasalazine and metformin. These drugs were added into the culture media of sFlt-1 overexpression organoids for a period of 7 days prior to experimental testing (**Fig. 4a**).

**Figure 4.**
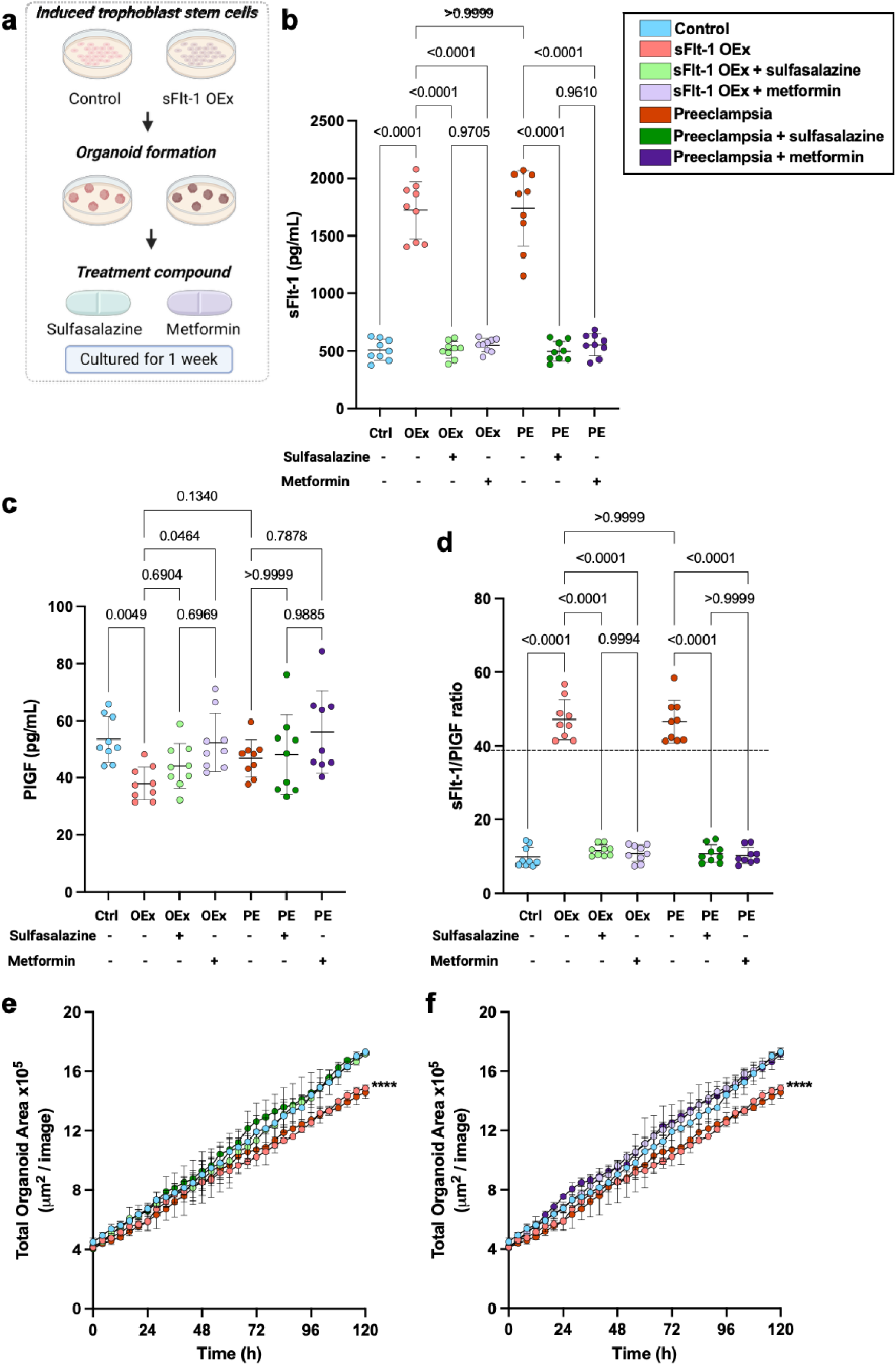
sFlt-overexpression trophoblast organoids are pharmacologically modulated by preeclampsia therapeutics. **a,** Workflow for experiments. Concentrations of angiogenic factors in culture media of control (Ctrl; blue), sFlt-1 OEx (OEx; coral), sFlt-1 OEx with sulfasalazine (light green), sFlt-1 OEx with metformin (light purple), primary preeclamptic (PE; red), primary preeclamptic with sulfasalazine (deep green) and primary preeclamptic with metformin (deep purple) trophoblast organoids, including **b,** sFlt-1 (pg/mL) **c,** PlGF (pg/mL) **d,** sFlt-1/PlGF ratio. The dashed line indicates the clinical cutoff for preeclampsia diagnosis (38). Organoid growth measured as total organoid area x10^5^ of **e,** sulfasalazine-treated and **f,** metformin-treated trophoblast organoids, using the Incucyte SX5 Live-Cell Imaging Platform with Incucyte Organoid Analysis Software. For **b-d**, data are presented as a scatter plot with interquartile range indicated and assessed using Welch’s ANOVA with multiple comparisons. For **e-f**, data are presented as mean ± standard deviation and assessed using a two-way ANOVA. n=9 trophoblast organoid replicates per group. *p < 0.05, **p < 0.01, ***p < 0.001. Source data file is provided.

Sulfasalazine is a sulfonamide prodrug which has anti-inflammatory and antioxidant properties. Whilst its most common use is as a disease-modifying anti-rheumatic drug in cases of bowel disease and intestinal inflammation, it is classified as safe to use in pregnancy [39]. Its effectiveness in human placental tissues from preeclamptic pregnancies has been demonstrated, and it reduces sFlt-1 and sEndoglin and upregulates PlGF secretion [22].

Metformin is a biguanide antihyperglycemic agent, commonly used as an antidiabetic agent to improve glucose tolerance. It is also classified as safe to use during pregnancy [39]. It has also shown to be effective in human preeclamptic placenta tissues, reducing secretion of sFlt-1 and sEndoglin, as well as VCAM1 [23].

Consistent with the model design, sFlt-1 overexpression organoids exhibited a marked increase in secreted sFlt-1 compared to control organoids, which was completely reversed by treatment with sulfasalazine and metformin (**Fig. 4b**). Primary preeclampsia organoids similarly exhibited elevated sFlt-1 concentrations, comparable to those observed in sFlt-1 overexpression organoids, which were significantly reduced by both sulfasalazine and metformin treatment. The modest reduction in PlGF levels observed in sFlt-1 overexpression organoids was completely reversed with sulfasalazine treatment and partially ameliorated with metformin treatment (**Fig. 4c**). The resultant sFlt-1/PlGF ratio (**Fig. 4d**) which is increased in sFlt-1 overexpression trophoblast organoids and primary preeclamptic organoids was completely reversed with sulfasalazine and metformin treatments. These findings demonstrate that the engineered organoids are responsive to pharmacological agents known to target pathways relevant to preeclampsia.

To determine whether these treatments impacted organoid growth, we quantified total organoid area over time. sFlt-1 overexpression and primary preeclamptic trophoblast organoids were significantly smaller and grew more slowly, and this trajectory was rescued by sulfasalazine treatment (**Fig. 4e**) as well as by metformin treatment (**Fig. 4f**) for both groups.

Together, these data show that sFlt-1 overexpression trophoblast organoids reproduce pathological features of preeclampsia and respond to clinically relevant therapeutics, supporting their utility as a human model for studying disease mechanisms and treatment responses.

### Conditioned media from sFlt-1 overexpression trophoblast organoids induces endothelial dysfunction

As endothelial dysfunction in the vessels and the heart is a central feature of preeclampsia pathology and a downstream consequence of excess circulating sFlt-1, we next sought to determine whether factors secreted by sFlt-1 overexpression trophoblast organoids were sufficient to impair endothelial function. To test this, conditioned media collected from control and sFlt-1 overexpression organoids was applied to primary human coronary artery endothelial cells (HCAECs) and primary human umbilical vein endothelial cells (HUVECs) in a tube formation assay (**Fig. 5a**).

**Figure 5.**
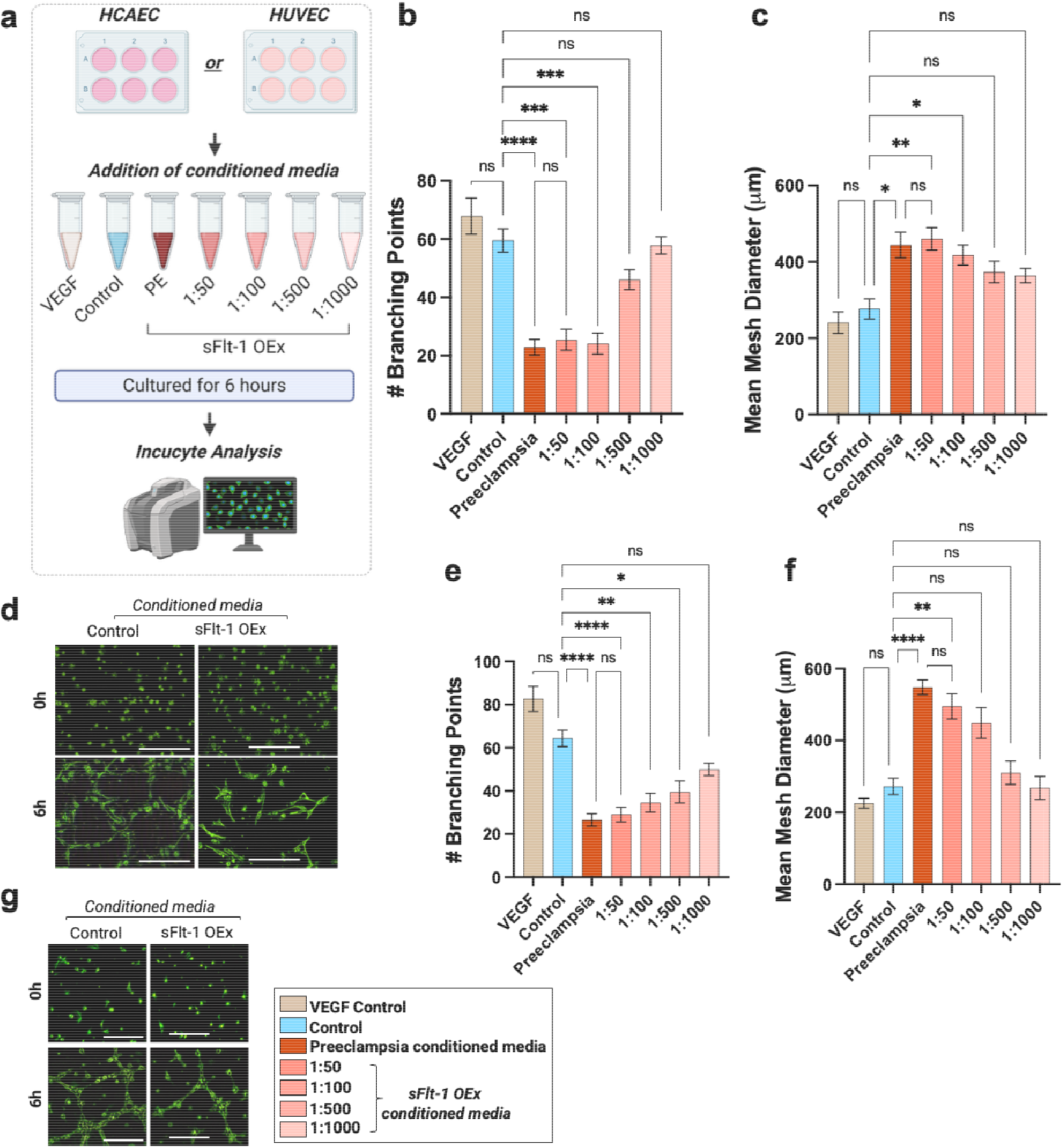
Conditioned media from sFlt-1 overexpression trophoblast organoids decreases endothelial tube formation. Quantification of endothelial tube formation by primary human coronary artery endothelial cells (HCAECs) or primary human umbilical vein endothelial cells (HUVECs) following exposure to VEGF (brown) or conditioned media collected from control (blue), primary preeclamptic or sFlt-1 overexpression (sFlt-1 OEx; coral) trophoblast organoids. **a,** Workflow for experiment. **b,** Number of branching points and **c,** mean mesh diameter of HCAECs were quantified following culture on synthetic hydrogel for 6 hours. **d,** Representative images of HCAEC tube formation following exposure to conditioned media from control and sFlt-1 OEx trophoblast organoids. **e,** Number of branching points and **f,** mean mesh diameter of HUVECs were quantified following culture on a synthetic hydrogel for 6 hours. **g,** Representative images of HUVEC tube formation following exposure to conditioned media from control and sFlt-1 OEx trophoblast organoids. Scale bar for **d,g,** = 200 μm. For **b-c**, data are presented as a histogram ± standard deviation and assessed using Welch’s ANOVA with multiple comparisons. n=9 replicates per group. * p < 0.05; ** p < 0.01; *** p < 0.001; **** p < 0.0001; ns = not significant (p ≥ 0.05). Source data file is provided.

Exposure of HCAECs to conditioned media from sFlt-1 overexpression organoids significantly impaired endothelial network formation in a dose-dependent manner. The number of branching points was reduced at higher concentrations of sFlt-1 overexpression organoid-conditioned media, with the greatest reduction observed at 1:50 (mean = 25.44) and 1:100 (mean = 24.11) dilutions compared to the control (mean = 59.56) and VEGF control (mean = 67.89) (**Fig. 5b**). Mean mesh diameter was also increased, reaching significance at the 1:50 (mean = 460.40 μm) and 1:100 dilutions (mean = 417.90 μm) compared to the control (mean = 276.61 μm) and VEGF control (mean = 240.74 μm) (**Fig. 5c**). There was no difference in number of branching points (mean = 22.89) nor mean mesh diameter (mean = 444.33 μm) in HCAECs exposed to conditioned media from primary preeclamptic organoids compared with sFlt-1 overexpression organoids.

The HCAECs response was independently recapitulated in HUVECs. Exposure to conditioned media from sFlt-1 overexpression organoids significantly impaired endothelial network formation compared with control-conditioned media, in a dose-dependent manner. HUVECs exposed to sFlt-1 overexpression organoid-conditioned media exhibited significantly fewer branching points at the 1:50 (mean = 29.11), 1:100 (mean = 34.56) and 1:500 (mean = 39.56) dilutions compared to the control (mean = 64.33) and VEGF control (mean = 82.67) (**Fig. 5e**). Mean mesh diameter was similarly increased at the 1:50 (mean = 495.00 μm) and 1:100 (mean = 448.44 μm) dilutions compared to the control (mean = 272.89 μm) and VEGF control (mean = 225.33 μm) (**Fig. 5f**), consistent with disruption of stable vascular network formation. There was no difference in number of branching points (mean = 26.44) nor mean mesh diameter (mean = 548.43 μm) in HUVECs exposed to conditioned media from primary preeclamptic organoids compared with sFlt-1 overexpression organoids.

Representative phase-contrast images further demonstrated altered endothelial morphology following exposure to conditioned media from sFlt-1 overexpression organoids (**Fig. 5d,g**). Whereas control-conditioned media supported formation of interconnected endothelial networks, conditioned media from sFlt-1 overexpression organoids resulted in fragmented structures with reduced branching complexity and impaired tube organisation.

These findings collectively demonstrate that the secretome of sFlt-1 overexpression trophoblast organoids is sufficient to induce endothelial dysfunction, recapitulating a key downstream pathological consequence of preeclampsia.

## Discussion

Here, we establish a genetically engineered trophoblast organoid model of preeclampsia generated independently of placental tissue access. Using a CRISPR Prime Integrase strategy, we engineered induced trophoblast stem cells to constitutively express the preeclampsia-associated sFlt-1 exon 15a isoform, generating a tractable human system that recapitulates molecular and functional features of disease. Significantly, overexpression of a single disease-associated sFlt-1 isoform was sufficient to induce broader changes in trophoblast gene expression, dysfunction and generates a secretome capable of impairing endothelial function.

A major advantage of this model is that it overcomes dependence on primary preeclamptic tissues. Access to diseased placental samples is inherently restricted to pregnancies already affected by pathology and generally occurs at delivery[40], limiting investigation of early disease processes and creating substantial barriers to reproducibility and scalability. In contrast, the present model can be generated reproducibly from induced trophoblast stem cells, providing an experimentally accessible platform for investigating preeclampsia biology independent of tissue availability. Transcriptomic analyses demonstrated that sFlt-1 overexpression organoids shifted toward the molecular space occupied by primary preeclamptic placentae, while maintaining human trophoblast context. This shift was accompanied by altered expression of genes implicated in preeclampsia pathology, including *FLT1*, *ENG*, *ET1* and *VCAM1*. Similarly, pathways including VEGF signalling, angiogenesis, vasoconstriction and vasculogenesis were enriched. These findings indicate that sustained sFlt-1 exon 15a expression induces coordinated transcriptional changes extending beyond the engineered FLT1 perturbation itself.

Consistent with the transcriptional shift, the engineered organoids reproduced multiple hallmarks of preeclampsia. Beyond elevated secretion of total sFlt-1 and the disease-associated exon 15a isoform, organoids exhibited reduced PlGF secretion and developed an sFlt-1/PlGF ratio comparable to primary preeclamptic trophoblast organoids and exceeding the ratio in maternal blood of 38 used for clinical decision making in hypertensive pregnant women[1]. Additional disease-associated phenotypes emerged spontaneously, including oxidative stress, elevated IL-6 and soluble endoglin secretion, impaired growth and altered transcriptional programs associated with angiogenesis and vascular regulation. The similarity between the engineered and primary preeclamptic organoids is particularly notable, with comparable sFlt-1/PlGF ratios, impaired growth and downstream endothelial effects despite their distinct origins.

Notably, these broader phenotypes could not be attributed simply to increased extracellular sFlt-1. Despite achieving similarly elevated concentrations of total sFlt-1, direct supplementation of recombinant human sFlt-1 to control organoids did not reproduce the reduction in PlGF or increases in IL-6 and soluble endoglin observed in the engineered organoids. This suggests that sustained trophoblast expression of the exon 15a-containing isoform initiates broader changes in trophoblast biology than does extracellular (rh) sFlt-1 exposure alone. This distinction is important for disease modelling, as recombinant sFlt-1 exposure may reproduce the extracellular anti-angiogenic environment without capturing the trophoblast-intrinsic consequences of sustained isoform production. The accompanying oxidative stress and transcriptional changes further suggest that sFlt-1 exon 15a overexpression participates in a broader trophoblast stress response, although the mechanisms linking its expression to these secondary phenotypes remain to be determined.

A further strength of this system is its utility for downstream functional studies. Conditioned media from sFlt-1 overexpression organoids impaired endothelial tube formation, demonstrating that the trophoblast secretome generated by the model is sufficient to induce endothelial dysfunction. This effect was reproduced in two primary endothelial cell types, HUVECs and HCAECs, was concentration-dependent, and was comparable to that induced by conditioned media from primary preeclamptic organoids. Inclusion of coronary artery endothelial cells extends these findings beyond the commonly used HUVEC model and is particularly relevant given the systemic vascular and cardiovascular manifestations of preeclampsia. Together, these experiments connect the engineered trophoblast phenotype with a key downstream functional consequence of disease and provide a platform for identifying the secreted factors responsible for endothelial dysfunction.

The model also demonstrated responsiveness to therapeutics previously reported to modulate placental dysfunction in preeclampsia. Treatment with sulfasalazine and metformin normalised angiogenic imbalance and restored trophoblast growth trajectories, supporting the biological relevance of the system. Both treatments reduced sFlt-1 and normalised the sFlt-1/PlGF ratio in engineered and primary preeclamptic organoids, while also rescuing impaired growth. These concordant responses provide pharmacological validation of the engineered model and support its potential for drug discovery. Candidate therapies could therefore be evaluated across multiple placental phenotypes and, through conditioned-media endothelial assays, for their capacity to improve downstream endothelial function. This is particularly valuable given that preeclampsia therapeutic development remains constrained by limited access to human placental tissue and the poor translational fidelity of animal models[41].

Several limitations warrant consideration. The present study used a single engineered background and focused specifically on overexpression of the exon 15a isoform, whereas preeclampsia represents a heterogeneous disorder with multiple placental and maternal contributors. The constitutive expression system also does not reproduce the endogenous regulation or temporal dynamics of *FLT1* alternative splicing during pregnancy, and therefore represents a model of sustained sFlt-1 exon 15a-driven trophoblast dysfunction rather than the full complexity preeclampsia. The model additionally lacks maternal immune, vascular and haemodynamic components that contribute to disease progression *in vivo*. Future work incorporating additional genetic backgrounds and multicellular co-culture systems may further expand its translational utility. The system could also be adapted to introduce additional genetic or molecular perturbations associated with distinct preeclampsia subtypes, providing a framework for modelling disease heterogeneity in a controlled human trophoblast background.

Together, these findings establish a human trophoblast organoid model of preeclampsia that can be generated independently of placental tissue access while recapitulating key molecular and functional features of disease. By linking sustained expression of a disease-associated sFlt-1 isoform with trophoblast dysfunction, an altered secretome and downstream endothelial impairment, this model provides a reproducible human system in which molecular drivers of placental dysfunction can be directly connected to their functional consequences. Its scalability and responsiveness to pharmacological intervention provide a foundation for mechanistic studies and therapeutic discovery in preeclampsia.

## Acknowledgements

We thank the women who consented to their placental tissue being used for this research. Figures were created using BioRender.com

## Author Contributions

A.L.A. designed the study, performed experiments, collected and analysed data, and wrote the manuscript. C.L. designed and generated the plasmids, contributed to experimental design and edited the manuscript. D.L.M.G. and A.M. performed experiments, collected and analysed data, and reviewed the manuscript. G.M. performed cell reprogramming, contributed to experimental design and reviewed the manuscript. L.P., F.A. and P.Q.T. contributed to experimental design and reviewed the manuscript. A.B. performed experiments and reviewed the manuscript. J.M.P. provided iTSCs and reviewed the manuscript. C.T.R. provided supervision, funding acquisition, review and editing of the manuscript. All authors contributed to the article and approved the submitted version.

---

**Supplementary Table 1.**
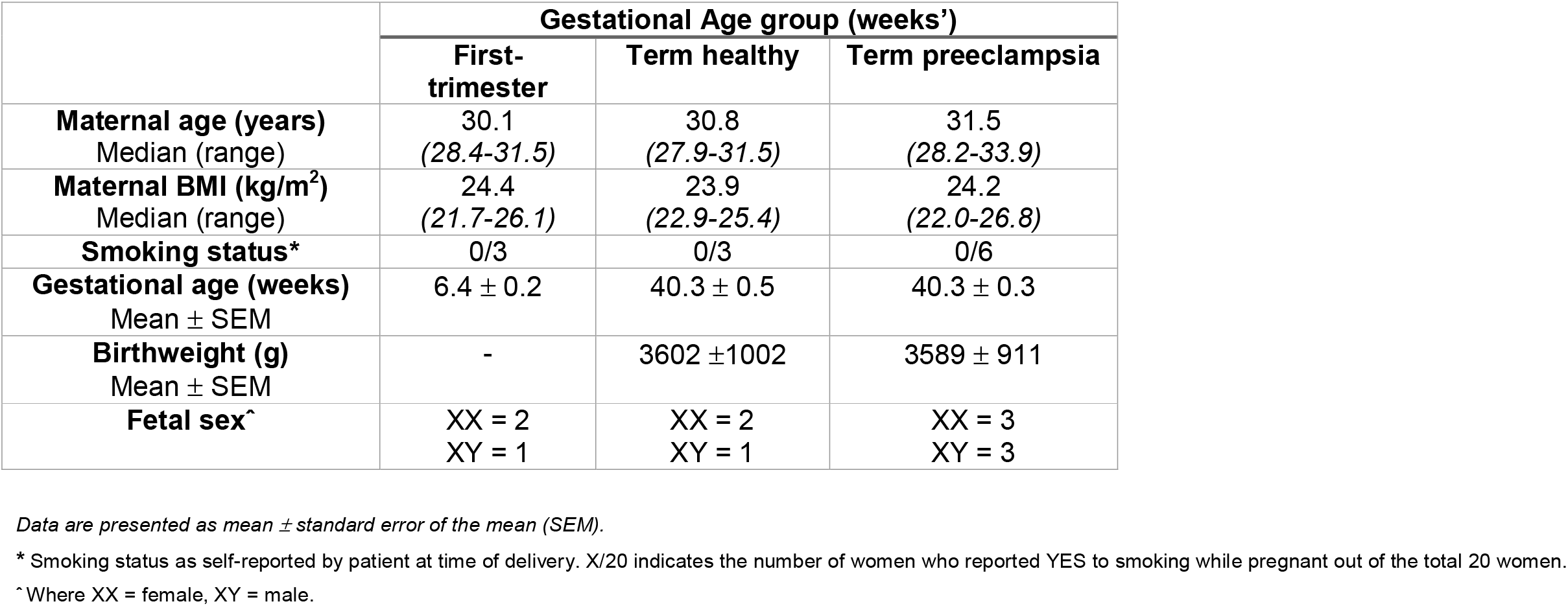
Characteristics of women who donated their placentae for this project.

|  | **Gestational Age group (weeks’)** |  |  |
|  | **First-trimester** | **Term healthy** | **Term preeclampsia** |
|---|---|---|---|
| **Maternal age (years)**<br>Median (range) | 30.1<br>*(28.4-31.5)* | 30.8<br>*(27.9-31.5)* | 31.5<br>*(28.2-33.9)* |
| **Maternal BMI (kg/m<sup>2</sup>)**<br>Median (range) | 24.4<br>*(21.7-26.1)* | 23.9<br>*(22.9-25.4)* | 24.2<br>*(22.0-26.8)* |
| **Smoking status\*** | 0/3 | 0/3 | 0/6 |
| **Gestational age (weeks)**<br>Mean ± SEM | 6.4 ± 0.2 | 40.3 ± 0.5 | 40.3 ± 0.3 |
| **Birthweight (g)**<br>Mean ± SEM | - | 3602 ±1002 | 3589 ± 911 |
| **Fetal sexˆ** | XX = 2<br>XY = 1 | XX = 2<br>XY = 1 | XX = 3<br>XY = 3 |

*Data are presented as mean ± standard error of the mean (SEM).*
\* Smoking status as self-reported by patient at time of delivery. X/20 indicates the number of women who reported YES to smoking while pregnant out of the total 20 women.
ˆ Where XX = female, XY = male.

